# *Atf-6* regulates lifespan through ER-mitochondrial calcium homeostasis

**DOI:** 10.1101/2020.01.14.906693

**Authors:** Kristopher Burkewitz, Sneha Dutta, Charlotte A. Kelley, Michael Steinbaugh, Erin J. Cram, William B. Mair

## Abstract

Functional crosstalk between organelles is critical for maintaining cellular homeostasis. Individually, dysfunction of both endoplasmic reticulum (ER) and mitochondria have been linked to cellular and organismal aging, but little is known about how mechanisms of inter-organelle communication might be targeted to extended longevity. The metazoan unfolded protein response (UPR) maintains ER health through a variety of mechanisms beyond its canonical role in proteostasis, including calcium storage and lipid metabolism. Here we provide evidence that in *C. elegans*, inhibition of the conserved UPR mediator, activating transcription factor (*atf*)*-6* increases lifespan via modulation of calcium homeostasis and signaling to the mitochondria. Loss of *atf-6* confers long life via downregulation of the ER calcium buffering protein, calreticulin. Function of the ER calcium release channel, the inositol triphosphate receptor (IP3R/*itr-1*), is required for *atf-6* mutant longevity while a gain-of-function IP3R/*itr-1* mutation is sufficient to extend lifespan. IP3R dysfunction leads to altered mitochondrial behavior and hyperfused morphology, which is sufficient to suppress long life in *atf-6* mutants. Highlighting a novel and direct role for this inter-organelle coordination of calcium in longevity, the mitochondrial calcium import channel, *mcu-1*, is also required for *atf-6* mutant longevity. Altogether this study reveals the importance of organellar coordination of calcium handling in determining the quality of aging, and highlights calcium homeostasis as a critical output for the UPR and *atf-6* in particular.

---

Aging is associated with failures in the ability to maintain homeostasis at the molecular and subcellular levels, and understanding how to prevent these age-related changes is essential to developing therapies for a variety of age-onset diseases. The ER acts as a hub of cellular metabolism and homeostasis through a variety of critical roles in the cell, which include housing the secretory pathway and associated proteostasis machineries, providing storage and timely release of cell calcium, and acting as the primary site of synthesis of triacylglycerols and membrane lipids^1^. Perturbations in any of these processes can trigger cellular dysfunction and lead to disease, necessitating a robust homeostatic network to maintain the health and function of the ER, the Unfolded Protein Response (UPR)^2^. As the name suggests, the UPR centers around its activation by unfolded proteins in the secretory pathway^2^. While activation of the UPR is most commonly associated with cytoprotection, especially as it relates to proteostasis, chronic UPR activation can also promote disease and pathology, in some cases limiting lifespan through its effects later in life^3,4^. Although the importance of the proteostasis outputs of the ER and UPR are well-established as critical for mediators of healthy aging^5^, how other diverse functions of the ER impact longevity are not well understood.

Three conserved branches mediate the UPR in metazoans, namely the inositol-requiring enzyme-1 (Ire1) branch, PKR-like endoplasmic reticulum kinase (PERK), and activating transcription factor 6 (*atf-6*). Previous studies of the UPR in mammalian models revealed that loss of Ire1 or PERK results in lethality or severe metabolic pathology, respectively, in mice, demonstrating critical roles for these branches in both organelle and organismal homeostasis^6,7^. In contrast, the first Atf6α knockout models surprisingly did not result in overt phenotypes^8,9^. Upon chronic, non-physiological dosing of tunicamycin, an inhibitor of protein glycosylation and maturation in the ER, however, Atf6α^-/-^ animals succumb to liver failure, revealing its importance in long-term adaptation to ER stress conditions^8,9^. Intriguingly, similar studies in *C. elegans* confirmed this hierarchical, relative importance of the three branches, revealing that *ire-1*/*xbp-1* and PERK/*pek-1* mutants exhibit the strongest phenotypes and are highly sensitive to tunicamycin, while *atf-6* mutants appear to be relatively unaffected by the toxin and present few baseline phenotypes^10–12^. Together these results suggest 1) that targeting *atf-6* may be more tolerable than the alternative UPR branches, and 2), *atf-6* may play a role in maintaining ER and cellular homeostasis beyond protein folding.

To define novel roles for ATF-6 in healthy aging we employed a *C. elegans* deletion mutant, *atf-6(ok551)*. First, we confirmed that *atf-6* mutants do not exhibit reduced survival in the presence of tunicamycin, an inhibitor of protein glycosylation and maturation in the ER lumen (Fig. 1A). Indicating that ATF-6 still functions to regulate some aspect of ER homeostasis in the nematode, loss of *atf-6* results in synthetic lethality when combined with loss of the *ire-1/xbp-1* pathway in *C. elegans*^*11*^. Furthermore, the ER residency and activation of mammalian Atf6 are accomplished through a transmembrane domain and lumenal Golgi trafficking signals, which are conserved in the nematode^11^.

**Figure 1.**
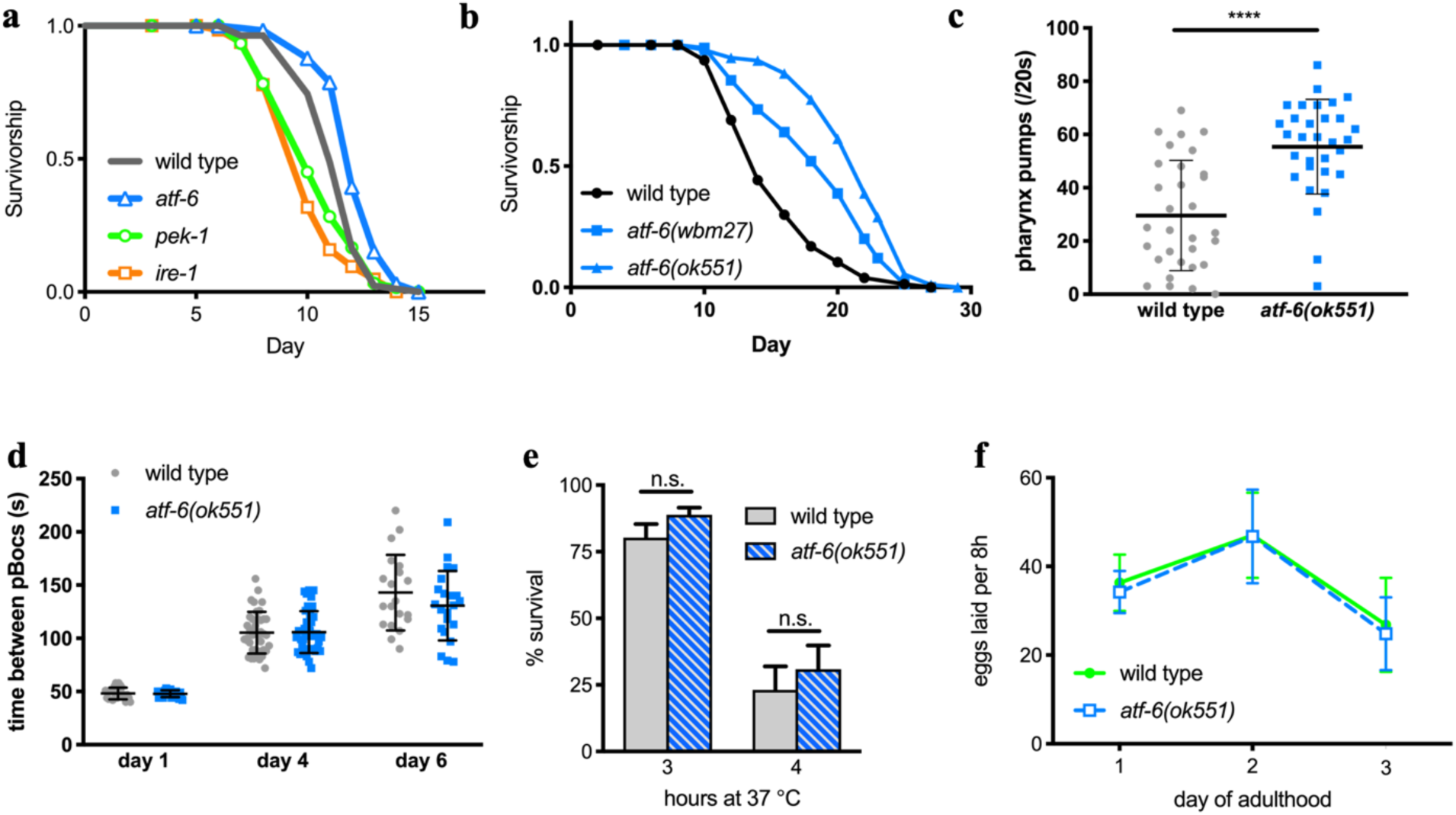
*Atf-6* mutants are not sensitive to proteotoxic stress and exhibit extended lifespan. a) Survival analysis of UPR mutant nematodes exposed chronically to 30 μg/mL tunicamycin starting on the first day of adulthood (n = 72 per condition). b) Lifespan analysis of *atf-6* mutants (n = 100 per condition). c) Pharyngeal pumping rates as a measure of healthspan in 7 day old nematodes (means +/- SD of n = 31 combined over 2 independent repeats). d) Period of the defecation motor program, an ultradian behavioral rhythm, in aging nematodes (n = 20-40 total periods measured from 4-5 animals on each day). e) Survival of worms exposed to high temperature for 3 and 4 hours (means + SD of 3 independent assays of 100 animals). f) Fecundity in *atf-6* mutants over the first 3 days of adulthood (means +/- SD of n = 5-19 worms per day).

To determine how *atf-6* impacts the aging process, we asked whether *atf-6* loss alters *C. elegans* lifespan. Counterintuitively for loss of a homeostatic regulator, we found that the *atf-6* deletion mutant was significantly long-lived (Fig. 1B), as had been suggested previously^13,14^. Confirming that this lifespan effect was due to loss-of-function of *atf-6*, we created a second null allele of *atf-6* through an early CRISPR/Cas9-generated frameshift, and this also extended lifespan (Fig. 1B). Furthermore, a subset of healthspan markers were improved in aged *atf-6* mutants, including pharyngeal pumping (Fig. 1C) but not rhythmicity of the defecation cycle (Fig. 1D), suggesting that normal *atf-6* function somehow promotes age-onset dysfunction and deterioration. We investigated whether loss of *atf-6* protected animals from heat, a more generalized form of proteotoxic stress that affects all intracellular compartments, and again found no differences from wild type animals (Fig. 1E). Due to the lack of sensitivity to the acute stress from either tunicamycin^8,11^ or heat treatment (Fig. 1A,E), we hypothesized that *atf-6* may instead play a role in basal ER functions, such as the biosynthetic demands associated with growth and reproduction. When we examined *atf-6* mutants for developmental or brood-size effects, however, we observed no developmental delay (data not shown) or defects in reproductive rate relative to wild type animals (Fig. 1F). Taken together, these findings suggest that in contrast to IRE1 and PERK, ATF-6 is pro-aging under basal conditions in *C. elegans*, providing an opportunity to discover novel longevity mechanisms downstream of this conserved transcription factor.

Given the lack of obvious phenotypes and putative mechanisms for *atf-6* longevity, we performed unbiased transcriptomic analyses to understand the physiological roles of *atf-6* in *C. elegans*. In agreement with previous analysis of *atf-6* dependent transcripts in early development^11^, we found that relatively few genes were differentially expressed (DE) in day-1 *atf-6(ok551)* adults when compared with wild-type N2 animals: only 26 genes were downregulated (Figure 2A, Table S1), while 101 genes were upregulated (Table S2) using cut-off criterion α < 0.01. Confirming previous studies in *C. elegans* and our hypothesis that *atf-6* may specialize in roles independent from the canonical UPR and proteostasis functions in *C. elegans*, we found virtually no signature markers of *ire-1/xbp-1* activation among these DE genes^10–12^ (Table S1-S2). This finding argues strongly against one possible model of *atf-6* longevity where compensatory activation of an alternative UPR branch is responsible for the lifespan extension. Because constitutive activation of the *ire-1/xbp-1* branch is sufficient to extend lifespan in *C. elegans*^*15*^, we attempted to further rule this compensatory model out by inhibiting proximal sensors of ER stress, *ire-1* and *pek-1*, via RNAi and performing lifespan analyses in *atf-6* mutants (Fig. S1). We found that neither *ire-1* nor *pek-1* is fully required for *atf-6* mutant longevity, though it is technically difficult to completely rule out contributions from the alternative branches considering the synthetic lethality of *ire-1*/*xbp-1* and *atf-6* mutants during development^11^.

**Figure 2.**
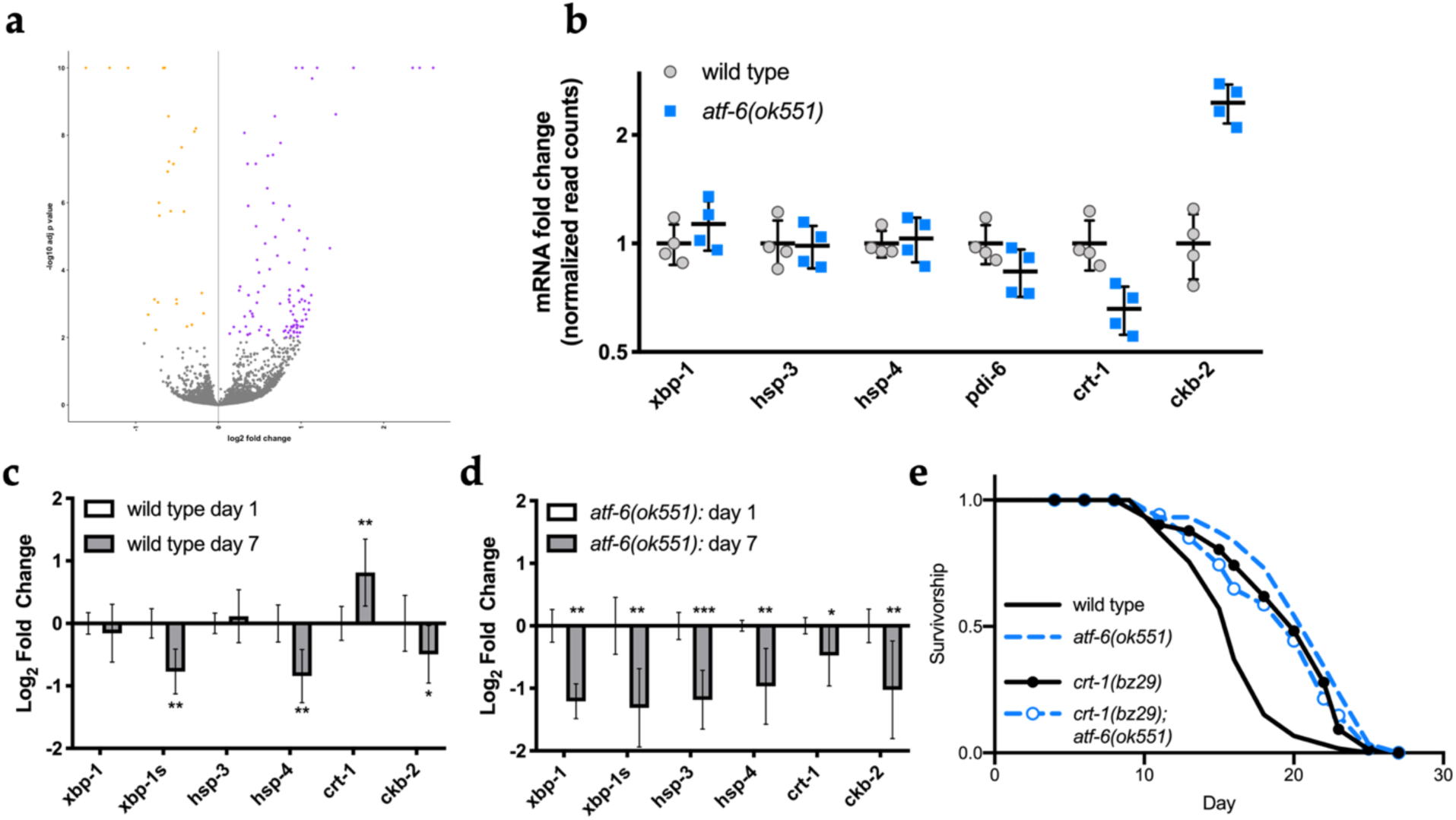
*Atf-6* regulates ER function and lifespan through its conserved target, calreticulin. a) Volcano plot of differentially expressed transcripts in *atf-6(ok551)* relative to N2. b) Relative transcript abundance of conserved UPR target genes in 1-day old *atf-6* mutants (mean +/- SD of RNA-Seq read counts normalized to wild type, n = 4). c, d) qRT-PCR analysis of changes in UPR transcripts in wild type worms between day 1 and day 7 of adulthood (c), contrasted with changes in UPR transcripts in aging *atf-6* mutants (d) (means +/- 95% confidence interval of 3 independent samples of ∼100 worms; *p<0.05, **p<0.01, ***p<0.001). e) Lifespan analysis of *atf-6(ok551)* and *crt-1(bz29)* mutants (n = 100 animals per curve).

The relatively small pool of DE genes suggests that *atf-6* dependent phenotypes arise from its regulation of single or small groups of transcriptional targets, as opposed to a large network effect. Among the most significantly affected transcripts, we found two genes previously implicated in ER stress responses: calreticulin/*crt-1*^*16*^ was downregulated in *atf-6* mutants, while choline kinase b-2/*ckb-2*^*17*^ was upregulated (Fig. 2B). Because *atf-6* mutants begin to show healthspan benefits after a week of adulthood, we reasoned that candidate aging factors may show the greatest differences in expression at that time point. We aged worms to 7 days and harvested RNA from wild type and *atf-6(ok551)* mutants. Strikingly, whereas *crt-1* levels are significantly upregulated with age in wild type animals (Fig. 2C), these transcripts are even further reduced with age in *atf-6* mutants (Fig. 2D), highlighting calreticulin as a potential mediator of *atf-6* longevity. If *atf-6* deletion is promoting lifespan through its effects on *crt-1*, we hypothesized loss of *crt-1* may be sufficient to reproduce the longevity phenotype. We employed a null mutant of calreticulin, *crt-1(bz29)*^*18*^, and consistent with our hypothesis, *crt-1(bz29)* mutants are long-lived to a similar extent as *atf-6(ok551)* (Fig. 2E). Together these results highlight calreticulin as a mediator of *atf-6* functions in *C. elegans* lifespan and are consistent with its evolutionary conservation as a direct target of Atf6 transcription in mammals^19^.

Calreticulin functions with calnexin as a quality control checkpoint during glycosylation of ER proteins, but plays a more outsized role as a calcium buffering protein, capable of binding and sequestering up to half of the total ER calcium pool^20^. Mirroring the lack of a proteostasis defect in *atf-6* mutants here, calnexin can effectively compensate for loss of calreticulin to maintain protein quality control in mammalian cells^21^. On the other hand, manipulating calreticulin levels causes overt alterations in calcium handling that ultimately drive changes in cell physiology^20^. Together these prior studies and our data suggest a model whereby *atf-6* maintains a primary role in the regulation of changes in ER calcium handling in parallel to *ire-1/xbp-1* and *pek-1* control of ER proteostasis. To test this concept, we placed synchronized L1 larval nematodes on bacterial lawns in the presence of thapsigargin, an inhibitor of ER calcium uptake, and measured their ability to grow and develop during chronic ER calcium stress. In contrast to what we observed with the protein-folding inhibitor tunicamycin^10–12^, growth of *atf-6* mutants is highly sensitive to thapsigargin and is reduced to the lowest levels among proximal UPR sensors (Fig. 3A).

**Figure 3.**
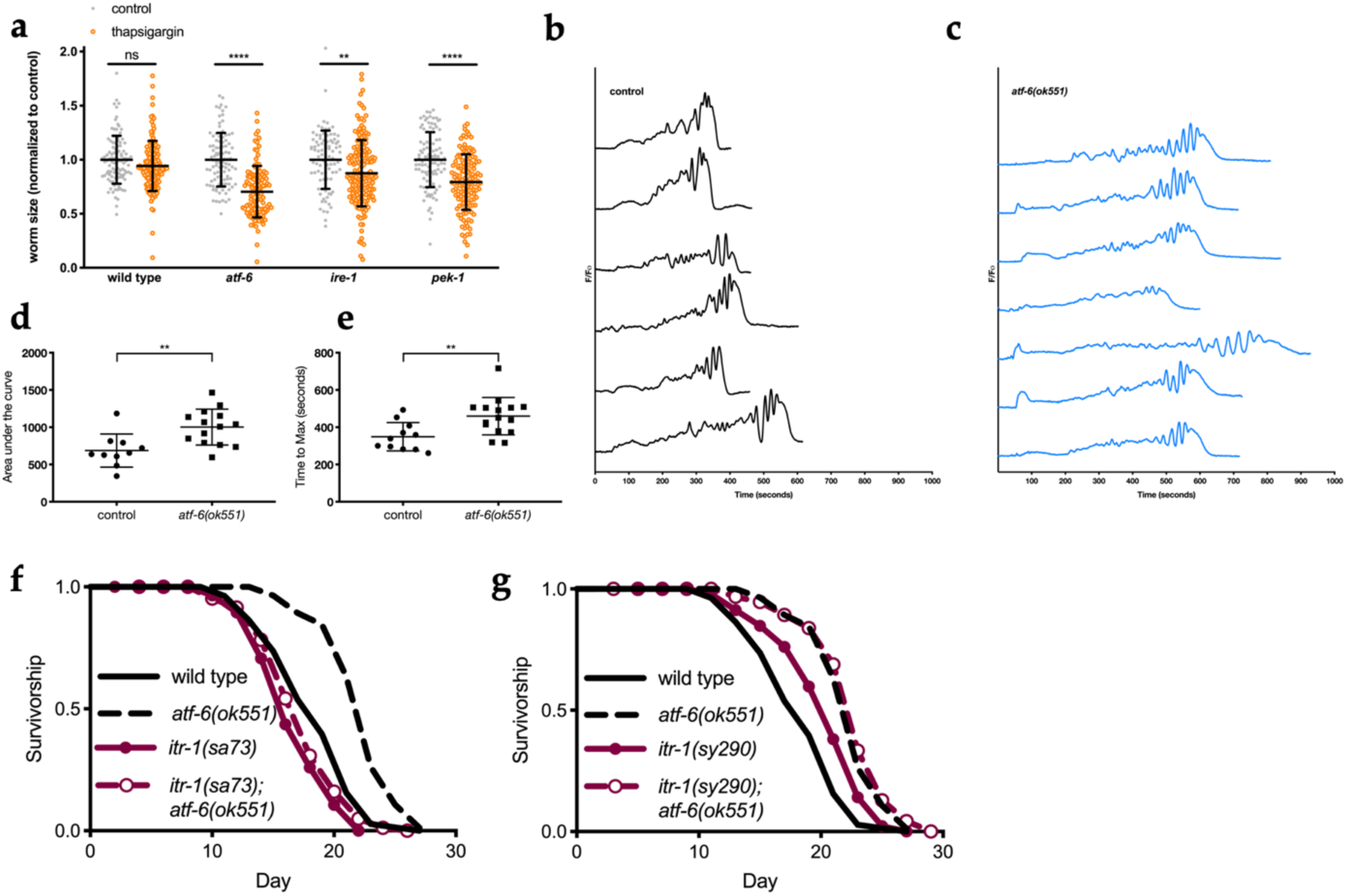
ER calcium flux functions downstream of *atf-6* in regulating lifespan. a) Relative growth of L1 larval worms exposed to 5 μg/mL of the SERCA inhibitor thapsigargin for 48 h (mean +/- SD of n = 88-164 worms combined over 2 independent trials). b, c) Representative traces of GCaMP3 imaging in the spermatheca of worms beginning with oocyte entry and ending with spermathecal contraction. d, e) Area under the curve (AUC) (d) and time-to-maximum (e) summary measurements of the GCaMP3 calcium traces recorded in the spermatheca of wild type and *atf-6* mutant worms (mean +/- SD of 11-16 animals combined over 2 repeats). f) Lifespan analysis of *atf-6* mutants in the temperature-sensitive IP3R/*itr-1(sa73)* background at the semi-permissive temperature of 20 C (n = 100 per condition). g) Lifespan analysis of the *atf-6* interaction with *itr-1(sy290)* gain-of-function mutation (n = 100 per condition). NS = p > 0.05, **p < 0.01, ***p < 0.001, ****p < 0.0001 by t-test.

These data suggest that *atf-6* may indeed play a role in maintaining ER calcium homeostasis, prompting us to determine whether there is evidence that *in vivo* calcium signaling is perturbed in these animals. One of the best-characterized calcium signaling paradigms in *C. elegans* is ovulation, where oocytes entering the smooth muscle-like spermatheca trigger oscillatory ER calcium release via the inositol triphosphate (IP_3_) receptor (IP3R), ultimately driving myosin-dependent contraction^22^. By expressing GCaMP3 in the spermatheca, we observed the cytosolic calcium oscillations and contraction events *in vivo* in *atf-6* mutants (Fig. 3B-C). These experiments revealed a number of aberrant phenotypes in *atf-6* mutants, including altered calcium oscillatory patterns, delayed calcium release and contraction, and greater total calcium release per contraction (Fig. 3B-E). Thus, *atf-6* appears to be a novel regulator of calcium homeostasis in *C. elegans*.

Due to the differences observed in the spermathecal calcium signaling of *atf-6* mutants and the central role for the IP3R in that process, we hypothesized ER calcium release via the IP3R may also play a role in the *atf-6* dependent lifespan effects. We crossed *atf-6* deletion mutants with both gain-of-function (*sy290*) and temperature-sensitive reduction-of-function (*sa73*) alleles of the sole *C. elegans* IP3R ortholog, *itr-1*. At the *sa73* semi-permissive temperature of 20 °C, *atf-6* longevity is fully suppressed in *itr-1(sa73)* mutants (Fig. 3F). Consistent with a model where enhanced IP3R-dependent calcium release promotes longevity, the gain-of-function allele, *itr-1(sy290)*, is sufficient to extend lifespan^23^, but its effects are not additive when crossed to the *atf-6* mutant (Fig. 3G). Together these results suggest a model where altered ER calcium capacity and release in *atf-6* mutants functions to extend lifespan.

We aimed next to better understand the mechanisms by which ER calcium release might be linked to longevity. Intriguingly, past studies have shown that calreticulin not only regulates ER calcium storage and release through the IP3R^20^, but also impacts mitochondrial calcium levels^24^. Furthermore, mild ER stress can result in organelle reorganization to increase communication between ER and mitochondria^25,26^. To determine how ER calcium efflux via *itr-1* can impact mitochondrial function and behavior, we performed *in vivo* imaging of mitochondrial networks in the nematode intestine. While young adult *atf-6* mutants themselves exhibited mild if any alterations in gross mitochondrial morphology, *itr-1(sa73)* mutants promoted dramatic reorganization of the networks, consistent with enhanced mitochondrial fusion (Fig. 4A-B). Because *itr-1(sa73)* both suppresses *atf-6* mediated longevity and causes mitochondrial networks to become hyperfused, we next asked whether this change in mitochondrial morphology was sufficient to block lifespan extension in this context. We fed *atf-6(ok551)* mutants dsRNA to inhibit the conserved fission factor, dynamin-related protein 1 (*drp-1)*, and this genetically imposed hyperfusion was indeed sufficient to block *atf-6* effects on lifespan (Fig. 4C). Conversely, enhancing mitochondrial fragmentation by feeding nematodes dsRNA to inhibit the ortholog of mammalian mitofusins, *fzo-1*, has no inhibitory effect on the longevity of *atf-6* mutants (Fig. 4D). Thus, reducing IP3R function causes mitochondrial hyperfusion, which is sufficient to suppress lifespan extension in *atf-6* mutants.

**Figure 4.**
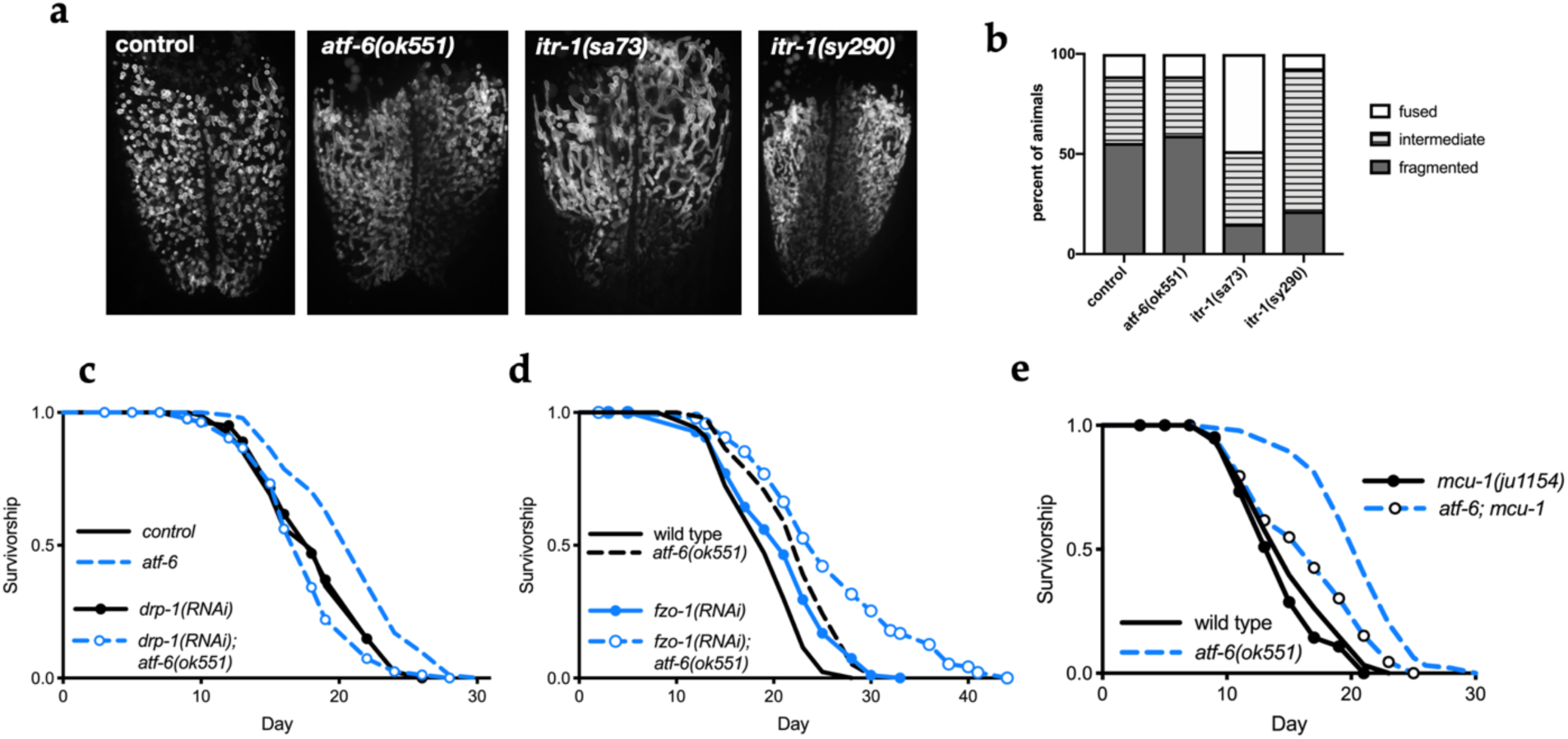
ER-mitochondrial communication via calcium regulates lifespan. a) Representative fluorescence Z-stack projections of intestinal mitochondrial networks via TOMM-20(1-49)::eGFP in day-1 adult animals. b) Quantification of mitochondrial morphology (n = 14-27 worms combined over 2 trials; P < 0.0001 comparing control vs. *itr-1(sa73)* and P = 0.001 comparing control vs. *itr-1(sy290)*). c, d) Lifespan analysis of worms fed dsRNA for the mitochondrial fission (*drp-1*, c) and fusion (*fzo-1*, d) machineries in control and *atf-6* mutant animals (n = 100 per condition). e) Lifespan analysis of *atf-6* mutants harboring a deletion in *mcu-1* to ablate acute mitochondrial calcium uptake (n = 100 per condition).

At ER-mitochondrial contact sites, calcium is directly transferred from the ER to the mitochondrial matrix via a series of transporters including the IP3R/*itr-1* on the ER surface, and voltage-gated anion channel (VDAC) and mitochondrial calcium uniporter (MCU)-1 in the mitochondrial outer and inner membranes, respectively^27^. In the matrix, calcium can stimulate metabolic and bioenergetic function through activation of one of several dehydrogenases^28,29^. Given that *atf-6* mutants show signs of a mild ER stress and that ER calcium efflux via *itr-1* is required for *atf-6* effects on lifespan, we hypothesized that ER-mitochondrial coordination of calcium homeostasis may be an important factor in this novel longevity mechanism. To test this, we crossed animals with deficient mitochondrial calcium import via MCU1/*mcu-1* deletion into the long-lived *atf-6(ok551)* background. Providing evidence that inter-organellar calcium flux is important in lifespan regulation, loss of the mitochondrial calcium importer, *mcu-1*, suppresses *atf-6* mediated lifespan extension (Fig. 4E).

In summary, we set out to identify the specialized roles *atf-6* plays in parallel to the PERK and Ire1/XBP-1 UPR pathways. We found in our characterization of the mutants that *atf-6* is a pro-aging factor, and deletion of this gene confers longevity in nematodes. Exploring the mechanism of *atf-6* longevity revealed ER calcium homeostasis as a critical downstream function, and highlighted calreticulin/*crt-1* as the proximal mediator of altered ER calcium storage and signaling. Furthermore, we demonstrated requirement and sufficiency for the highly conserved ER calcium release channel, the IP3R/*itr-1* in this pathway. Finally, in exploring the downstream mechanisms of longevity caused by modulation of ER calcium release, we discovered a key role for the mitochondrial uptake of calcium via *mcu-1*. These findings reveal the critical importance of non-canonical UPR outputs, namely ER calcium handling, in the aging process. Furthermore, we identified ER-mitochondrial coordination of calcium as a novel mechanism in lifespan regulation, opening exciting future avenues in understanding how calcium dyshomeostasis is not simply a marker for age-related disease, but a hallmark and driver of the aging process itself.

---

**Figure S1.**
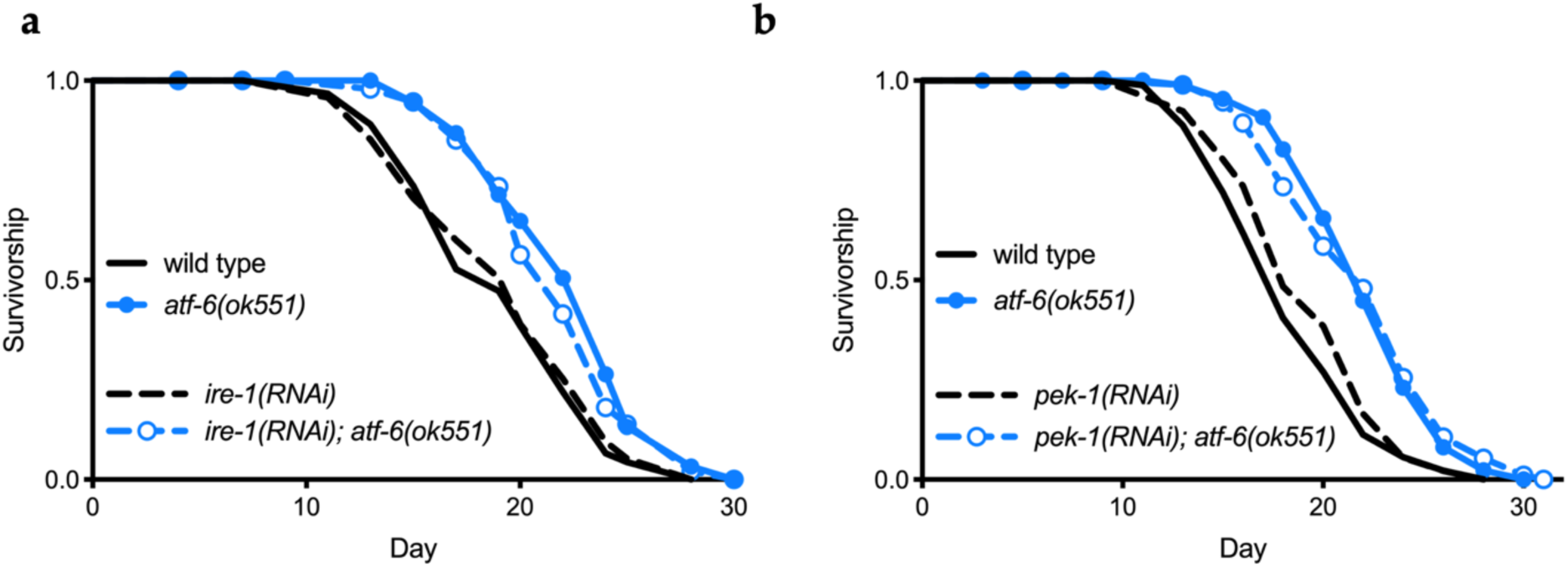
Longevity in *atf-6* mutants is parallel to the alternative UPR branches. a) Lifespan analysis of worms fed *ire-1* dsRNA starting on day 1 of adulthood. b) Lifespan analysis of worms fed *pek-1* dsRNA from hatch. Starting n = 100 animals per condition.

Table S1. Transcripts downregulated in *atf-6(ok551)* mutants on the first day of adulthood.

Table S2. Transcripts upregulated in *atf-6(ok551)* mutants on the first day of adulthood.

Table S3. Summary lifespan statistics.

## Materials and Methods

### *C. elegans* strains and husbandry

N2 wild-type, RB772 [*atf-6(ok551)*], SJ30 [*ire-1(zc14); zcIs4*], ZB1028 [*crt-1(bz29)*], CZ19982 [*mcu-1(ju1154)*], JT73 [*itr-1(sa73)*], RB545 [*pek-1(ok275)*], PS1631 [*itr-1(sy290); dpy-20(e1282)*] *C. elegans* strains were obtained from the *Caenorhabditis* Genetic Center, which is funded by NIH Office of Research Infrastructure Programs (P40 OD010440). High-throughput deletion strains (e.g., RB772 and RB545) were outcrossed 4-8 times with N2 before use, and PS1631 was backcrossed to N2 to isolate *itr-1(sy290)* from the *dpy-20* co-marker. Additional strains include WBM1093; fil-1p::GCaMP3, WBM926; WBM947; WBM973; WBM974; WBM1046; WBM1103; WBM1104; WBM1134; WBM1158. Worms were grown and maintained on standard nematode growth media (NGM) seeded with *E. coli* (OP50-1). *E. coli* bacteria was cultured overnight in LB at 37°C, after which 100 μl of liquid culture was seeded on plates to grow for 2 days at room temperature. RNAi experiments alternatively employed *E. coli* (HT115) from the Ahringer library (Source Bioscience) expressing dsRNA against the gene noted or an empty vector control. Experiments with HT115 were performed identically except LB and NGM contained 100 μg ml^-1^ Carbenicillin, and dsRNA expression was induced by addition of 100 μl IPTG (100 mM) at least 2 hours before worms were introduced to the plates.

### Survival Analyses

Lifespan experiments were performed on standard nematode growth media plates at 20 °C. Worms were synchronized by picking 1 day-old adult worms onto plates for timed egg lays, after which the adult worms were removed. When worms reached late L4/young adulthood, 100 worms were transferred to fresh plates at 10-25 worms per plate and this was considered time = 0. Worms were transferred to fresh bacterial lawns every other day to separate from progeny until the first deaths (10-14 d). Survival was scored every 1-2 days and a worm was deemed dead when unresponsive to 3 taps on the head and tail. Worms were censored due to contamination on the plate, worms leaving the agar media, progeny hatching inside the adult, or loss of vulval integrity during reproduction. To test survival on tunicamycin, toxin in DMSO (EMD Millipore) or DMSO control was added to NGM to a final concentration of 30 μg/mL before pouring 6-well plates. 10 μL of OP50-1 culture was spotted in each well 1 day before worms. Twelve day-1 adult worms were picked into each well for a total of 72 worms per sample. Worms were scored as above. Animals on tunicamycin plates were not transferred to separate progeny, as this concentration of tunicamycin effectively sterilizes the adults. Statistical significance for survival assays was determined through Mantel-Cox Log-rank (GraphPad Prism). Summary statistics for all survival experiments are presented in Table S3.

### Healthspan Assays

Worms were synchronized by egg lay and maintained as normal on OP50-1 lawns. After day-1 measurements, worms were transferred to fresh lawns at least every other day to separate progeny. To measure the oscillatory period of the defecation motor program, the timings of 8 consecutive pBocs were scored for 4-5 undisturbed worms of each genotype; after day 6 pBocs become severely irregular and each worm was measured for 20 minutes. Pharyngeal pumping was counted manually over 20 s intervals in undisturbed animals within the OP50-1 lawn. Worms which left the lawn during either assay were not recorded. Data were analyzed with t-tests (GraphPad Prism).

### Heat Stress

Worms synchronized by egg-lay were allowed to develop at 20 °C for ∼72 hours to adulthood. Once adults, 25 worms from each genotype were transferred to each of 4 fresh NGM plates with small OP50-1 lawns, for a total of 100 animals. Alternating by genotype, plates were arranged in a single layer in a 37 °C incubator until the respective timepoint, at which point plates were removed and returned to 20 °C to recover overnight. Survival was scored 24 h after stress by responsiveness to touch.

### Thapsigargin Stress

To test for growth during ER calcium stress, OP50-1 lawns were grown for 1 day before spotting thapsigargin in DMSO (Sigma) or DMSO control directly on to lawns to reach a final concentration of 5 μg/mL in the plate. Thapsigargin was given 24 hours to diffuse into the NGM before ∼30-60 synchronized L1s were introduced. After 48 hours, worms were washed off plates in M9 buffer, anesthetized in 2 mM sodium azide (Sigma), returned to unseeded NGM plates, and imaged at 4x magnification with a Zeiss Imager.M2 microscope. A binary mask was applied to images and the area of each worm was recorded (Image J). Data were analyzed with t-tests (GraphPad Prism).

### Transcriptomic Analysis

In each of 4 independent biological replicates, roughly 1000 hypochlorite-synchronized L1s were washed onto OP50-1 lawns and allowed to develop for 72 h to adulthood, before washing in M9 to remove bacteria, resuspending in Qiazol (Qiagen) and snap-freezing in liquid nitrogen. Total RNA was isolated after disrupting worms with 5 freeze-thaws with Qiazol extraction and RNeasy mini kit (Qiagen). cDNA libraries were prepared through TruSeq Stranded mRNA kit (Illumina) and sequenced with an Illumina NextSeq500 over 75 cycles by the Dana Farber Molecular Biology Core. Reads were analyzed through the bcbio pipeline and bcbioRNASeq R package (http://bioinformatics.sph.harvard.edu/bcbioRNASeq). Quality control with MultiQC was performed in parallel to differential expression with counts aligned to WBcel235 using STAR/featureCounts, revealing mapping rates >95% for ∼14-25 million total reads per replicate. Differential expression analysis was performed with DESeq2 via the bcbioRNASeq R package with counts generated by salmon (https://combine-lab.github.io/salmon). Significance was defined with adjusted P value (alpha; 1% false discovery rate) cutoff of 0.01.

### Gene Expression Assays

Total RNA was isolated from ∼100 L4 stage animals. RNA was extracted using Qiazol reagent (Qiagen) then column purified by RNeasy micro kit (Qiagen). cDNA was generated using SuperScript VILO master mix (Invitrogen). Taqman real-time qPCR experiments were performed on a StepOne Plus instrument (Applied Biosystems) following the manufacturer’s instructions. Data were analyzed with the comparative ΔΔ*C*t method using *Y45F10D*.*4* as endogenous control. ExpressionSuite software (Applied Biosystems) was used to generate average fold-change and 95% confidence intervals relative to day 1 adults and t-test for statistical significance.

### Mitochondrial morphology

Synchronized populations were grown to day-1 adults at 20 °C, and picked unanesthetized onto 10% agarose pads with 0.05 μm Polybead microspheres (Polysciences) for immobilization. Imaging was performed on a Ti2 CSU-W1 confocal microscope (Nikon) with 488-nm illumination of eGFP through a Plan-Apochromat 100x/1.45 objective. Qualitative assessment of mitochondrial morphology was made by blinded analysis, scoring worms based on three categories: tubular (interconnected mitochondrial network), intermediate (combination of interconnected network and isolated smaller mitochondria) or fragmented (mostly fragmented mitochondria). Differences between strains were tested for significance with Chi-square analysis (GraphPad Prism).

### Calcium Imaging

Partially synchronized populations were obtained by egg prep and animals were grown at 23°C for ∼54 h, around the time of the first ovulation. Live animals were immobilized with 0.01% tetramisole and 0.1% tricaine in M9 buffer and mounted on 2% agarose pads or with 0.05 μm Polybead microspheres (Polysciences) diluted 1:2 in water and mounted on 5% agarose pads. Confocal microscopy was performed on an LSM 710 confocal microscope (Zeiss) equipped with Zen software (Zeiss) using a Plan-Apochromat 63×/1.40 oil DIC M27 objective. A 488-nm laser was used for GCaMP3. Live animals were imaged for ∼30 min total.

### Contributions

K.B. and W.B.M. designed the study. K.B. designed and performed the majority of experiments. S.D. performed repeats of lifespan experiments. C.A.K. and E.J.C. designed and performed calcium imaging experiments. M.S. performed the RNA-sequencing analysis. The manuscript was written by K.B. with edits and feedback from all authors.

## References

1. Bravo, R. et al. Endoplasmic reticulum and the unfolded protein response: dynamics and metabolic integration. Int. Rev. Cell Mol. Biol. 301, 215–290 (2013).

2. Ron, D. & Walter, P. Signal integration in the endoplasmic reticulum unfolded protein response. Nat. Rev. Mol. Cell Biol. 8, 519–529 (2007).

3. Wang, L., Ryoo, H. D., Qi, Y. & Jasper, H. PERK Limits Drosophila Lifespan by Promoting Intestinal Stem Cell Proliferation in Response to ER Stress. PLoS Genet. 11, e1005220 (2015).

4. Hotamisligil, G. S. Endoplasmic reticulum stress and the inflammatory basis of metabolic disease. Cell 140, 900–917 (2010).

5. Frakes, A. E. & Dillin, A. The UPR(ER): Sensor and Coordinator of Organismal Homeostasis. Mol. Cell 66, 761–771 (2017).

6. Zhang, K. et al. The unfolded protein response sensor IRE1alpha is required at 2 distinct steps in B cell lymphopoiesis. J. Clin. Invest. 115, 268–281 (2005).

7. Zhang, P. et al. The PERK eukaryotic initiation factor 2 alpha kinase is required for the development of the skeletal system, postnatal growth, and the function and viability of the pancreas. Mol. Cell. Biol. 22, 3864–3874 (2002).

8. Wu, J. et al. ATF6alpha optimizes long-term endoplasmic reticulum function to protect cells from chronic stress. Dev. Cell 13, 351–364 (2007).

9. Yamamoto, K. et al. Transcriptional induction of mammalian ER quality control proteins is mediated by single or combined action of ATF6alpha and XBP1. Dev. Cell 13, 365–376 (2007).

10. Springer, W., Hoppe, T., Schmidt, E. & Baumeister, R. A Caenorhabditis elegans Parkin mutant with altered solubility couples alpha-synuclein aggregation to proteotoxic stress. Hum. Mol. Genet. 14, 3407–3423 (2005).

11. Shen, X., Ellis, R. E., Sakaki, K. & Kaufman, R. J. Genetic interactions due to constitutive and inducible gene regulation mediated by the unfolded protein response in C. elegans. PLoS Genet. 1, e37 (2005).

12. Bischof, L. J. et al. Activation of the unfolded protein response is required for defenses against bacterial pore-forming toxin in vivo. PLoS Pathog. 4, e1000176 (2008).

13. Wang, N. et al. miR-124/ATF-6, a novel lifespan extension pathway of Astragalus polysaccharide in Caenorhabditis elegans. J. Cell. Biochem. 116, 242–251 (2015).

14. Henis-Korenblit, S. et al. Insulin/IGF-1 signaling mutants reprogram ER stress response regulators to promote longevity. Proc. Natl. Acad. Sci. U. S. A. 107, 9730–9735 (2010).

15. Taylor, R. C. & Dillin, A. XBP-1 is a cell-nonautonomous regulator of stress resistance and longevity. Cell 153, 1435–1447 (2013).

16. Park, B. J. et al. Calreticulin, a calcium-binding molecular chaperone, is required for stress response and fertility in Caenorhabditis elegans. Mol. Biol. Cell 12, 2835–2845 (2001).

17. Caruso, M.-E. et al. GTPase-mediated regulation of the unfolded protein response in Caenorhabditis elegans is dependent on the AAA+ ATPase CDC-48. Mol. Cell. Biol. 28, 4261–4274 (2008).

18. Xu, K., Tavernarakis, N. & Driscoll, M. Necrotic cell death in C. elegans requires the function of calreticulin and regulators of Ca(2+) release from the endoplasmic reticulum. Neuron 31, 957–971 (2001).

19. Yoshida, H., Haze, K., Yanagi, H., Yura, T. & Mori, K. Identification of the cis-acting endoplasmic reticulum stress response element responsible for transcriptional induction of mammalian glucose-regulated proteins. Involvement of basic leucine zipper transcription factors. J. Biol. Chem. 273, 33741–33749 (1998).

20. Michalak, M., Groenendyk, J., Szabo, E., Gold, L. I. & Opas, M. Calreticulin, a multi-process calcium-buffering chaperone of the endoplasmic reticulum. Biochem. J 417, 651–666 (2009).

21. Molinari, M. et al. Contrasting functions of calreticulin and calnexin in glycoprotein folding and ER quality control. Mol. Cell 13, 125–135 (2004).

22. Kovacevic, I., Orozco, J. M. & Cram, E. J. Filamin and phospholipase C-ε are required for calcium signaling in the Caenorhabditis elegans spermatheca. PLoS Genet. 9, e1003510 (2013).

23. Iwasa, H., Yu, S., Xue, J. & Driscoll, M. Novel EGF pathway regulators modulate C. elegans healthspan and lifespan via EGF receptor, PLC-gamma, and IP3R activation. Aging Cell 9, 490–505 (2010).

24. Arnaudeau, S. et al. Calreticulin differentially modulates calcium uptake and release in the endoplasmic reticulum and mitochondria. J. Biol. Chem. 277, 46696–46705 (2002).

25. Bravo, R. et al. Increased ER-mitochondrial coupling promotes mitochondrial respiration and bioenergetics during early phases of ER stress. J. Cell Sci. 124, 2143–2152 (2011).

26. Arruda, A. P. et al. Chronic enrichment of hepatic endoplasmic reticulum-mitochondria contact leads to mitochondrial dysfunction in obesity. Nat. Med. 20, 1427–1435 (2014).

27. Szabadkai, G. et al. Chaperone-mediated coupling of endoplasmic reticulum and mitochondrial Ca2+ channels. J. Cell Biol. 175, 901–911 (2006).

28. Denton, R. M. Regulation of mitochondrial dehydrogenases by calcium ions. Biochim. Biophys. Acta 1787, 1309–1316 (2009).

29. Griffiths, E. J. & Rutter, G. A. Mitochondrial calcium as a key regulator of mitochondrial ATP production in mammalian cells. Biochim. Biophys. Acta 1787, 1324–1333 (2009).

